# Secondary microglia formation center in the human fetal brain

**DOI:** 10.1101/2024.02.29.582659

**Authors:** Chenyun Song, Xinyu Chen, Rong Ji, Yang Liu, Ling Zhang, Lu Gao, Qizhi He, Lixiang Ma, Hexige Saiyin

**Affiliations:** Department of Anatomy and Histology & Embryology, School of Basic Medical Sciences, Fudan University, Shanghai 200032; Department of Neurology, Huadong Hospital, Fudan University, Shanghai, P.R. China Department of Neurology, Huadong Hospital, Fudan University, Shanghai, P.R. China; Department of Developmental Biology, Washington University School of Medicine, St Louis, Missouri, United States of America (Current); Department of Pathology, Shanghai Key Laboratory of Maternal-Fetal Medicine, Shanghai Institute of Maternal-Fetal Medicine and Gynecologic Oncology, Shanghai First Maternity and Infant Hospital, School of Medicine, Tongji University, Shanghai 201204, P.R. China; State Key Laboratory of Genetic Engineering, School of Life Sciences, Fudan University, Shanghai 200433

**Author notes:** The authors equally contributed to this study (Chenyun Song, Xinyu Chen, Rong Ji).

## Abstract

Yolk sac-derived microglia migrate and populate the brain during development, constituting 10−15% of the total brain cells. The human brain is the largest and most complex brain with the highest cognitive capacity among all species. Therefore, the limitations of rodent brain studies in interpreting the human brain are evident. By co-immunostaining microglia in 50 µm fetal brain sections from 7.5 to 16 gestational weeks (gw) and combining high-resolution scanning, we identified a highly proliferative microglia aggregate (0.108−2.129 mm^2^) that expanded in Down’s Syndrome fetal brain (4.168 mm^2^) and was located near the ganglion eminence, in which Ki67^+^ microglia accounted for 23.4% of total microglia compared to 6.3% in other brain regions. The microglia in the aggregates lack phagocytic bulbs, membrane ruffles, and long/branching processes compared to microglia in other brain regions. Introducing human microglia into cortical organoids, but not macrophages, replicated proliferative microglial aggregates on the brain organoid surface and sufficiently penetrated deeper regions of the cortical organoids. Penetrating microglia display phagocytic capacity, enhance immunity, and accelerate the maturation of brain organoids. The large proliferative microglial aggregate may be a unique secondary microglial formation center in the human fetal brain to compensate for the enormous microglial demands during brain expansion.

## Introduction

Microglia are highly dynamic and versatile resident phagocytes that originate in the yolk sac (YS) ^1–5^. Microglia migrate from YS to the fetal brain around 4.5−5.5 gw, disproportionally reaching different brain regions and finally accounting for 10−15% of total brain cells^6, 7^. Functionally, microglia phagocytize effete cells, aberrant projections, repair neuronal wiring, prune excessive/abnormal synapses, and secrete cytokines to maintain brain homeostasis^8–10^. The size and complexity of the human brain are much greater than those of other lab species^7^. Furthermore, human microglia are more heterogeneous than those of other species^11^. However, it remains unclear how microglia from the YS fully populate the rapidly expanding fetal brain in a human-specific manner.

The human fetal brain has a large outer subventricular zone (OSVZ) in subventricular zone (SVZ), contributing to the large size and complexity of the human cortex^12^. The OSVZ forms at approximately13 gw, expands dramatically after its formation, and provides a scaffold for migrating cells^12, 13^. YS-derived microglia enter the fetal brain before generating astrocytes and oligodendrocytes and form OSVZ. Microglia reach the brain through circulation-dependent or circulation-independent routes in mice or zebrafish^14,15^. Microglia are amoeboid in early fetal stages and traverse the arachnoid membrane, brain ventricles, and choroid plexus to the developing cortex^16^. The shape change of the microglia in the parenchyma defines their orientation of the microglia^17^. After arriving at their destination, the microglia differentiate and mature. Emerging evidence reveals that microglial dysfunction is associated with a wide range of diseases, including obsessive-compulsive disorder, Rett syndrome, and neurodegenerative disorders such as Alzheimer’s disease (AD), Parkinson’s disease (PD)^18^ and Huntington’s disease (HD)^19^. Microglia are involved in moderate inflammatory reactions that engulf debris or protein deposits in a reasonable timeframe, and microglia involved in an excessive and sustained inflammatory response result in detrimental neuronal damage and loss of synapses^20, 21^.

Brain organoids derived from human induced pluripotent stem cells (hiPSCs) robustly recapitulate the histological and anatomical features of the early human fetal brain, especially the OSVZ, and provide a tangible tool for studying human-specific characteristics in developing brains^22–24^. However, most brain organoids lack microglia of the yolk (a YS origin)^24^. The introduction of human microglia into human brain organoids partially mimics microglial physiological roles and anatomical signatures in the human fetal brain and improves brain organoid maturation^25–27^. Therefore, the introduction of microglia into human brain organoids through different routes may reveal a human-specific evolution or populate the dynamics of microglia in the fetal brain.

Herein, we combined multiple antibodies’ immunostaining with state-of-the-art high-resolution scanning of sections 50 µm in thickness to see the morphological evolution and dynamics of human microglia in the human fetal brain. We also introduced microglia into human brain organoids to replicate the microglial dynamics and evolution in the fetal brain. Our study identified a microglial formation center in the fetal brain that paralleled the time-lapse of OSVZ formation and expanded dramatically in the Down’s syndrome (DS) fetal brain. The introduction of microglia into brain organoids replicated the center of microglia formation.

## Results

### Migrating microglia monitor and be scaffolded by radial glia in early fetal brain

The diameter of human microglia is approximately 50 µm^28^. Therefore, immunostaining in thin brain sections (<20 µm in thickness) makes it impossible to visualize intact human microglia. To see the spatiotemporal status of intact microglia in the human fetal brain, we used IBA-1, CD34, and Ki67 or IBA-1, Ki67, and vimentin antibodies immunostaining in 50 µm human fetal brain sections (7.5 −16 gw), and scanned the thick sections using high-resolution confocal microscopy, including structural illumination microscopy (SIM). Similarly to other findings^6, 29, 30^, microglia were amoeboid and rare at the early brain stage with 7.5 gw and increased following an increase in gw **(Fig.S1A, B)**; a limited number of microglia with amoeboid or bipolar morphology were distributed in the neuroepithelium of 7.5 gw, and few microglia were present in the ventricles **(Fig. S1A)**. Microglia processes became more complicated and branched, and the morphological complexity scores increased from 7.5 gw to 15 gw **(Fig. S1C**; **Fig. 1D)**.

**Fig. 1.**
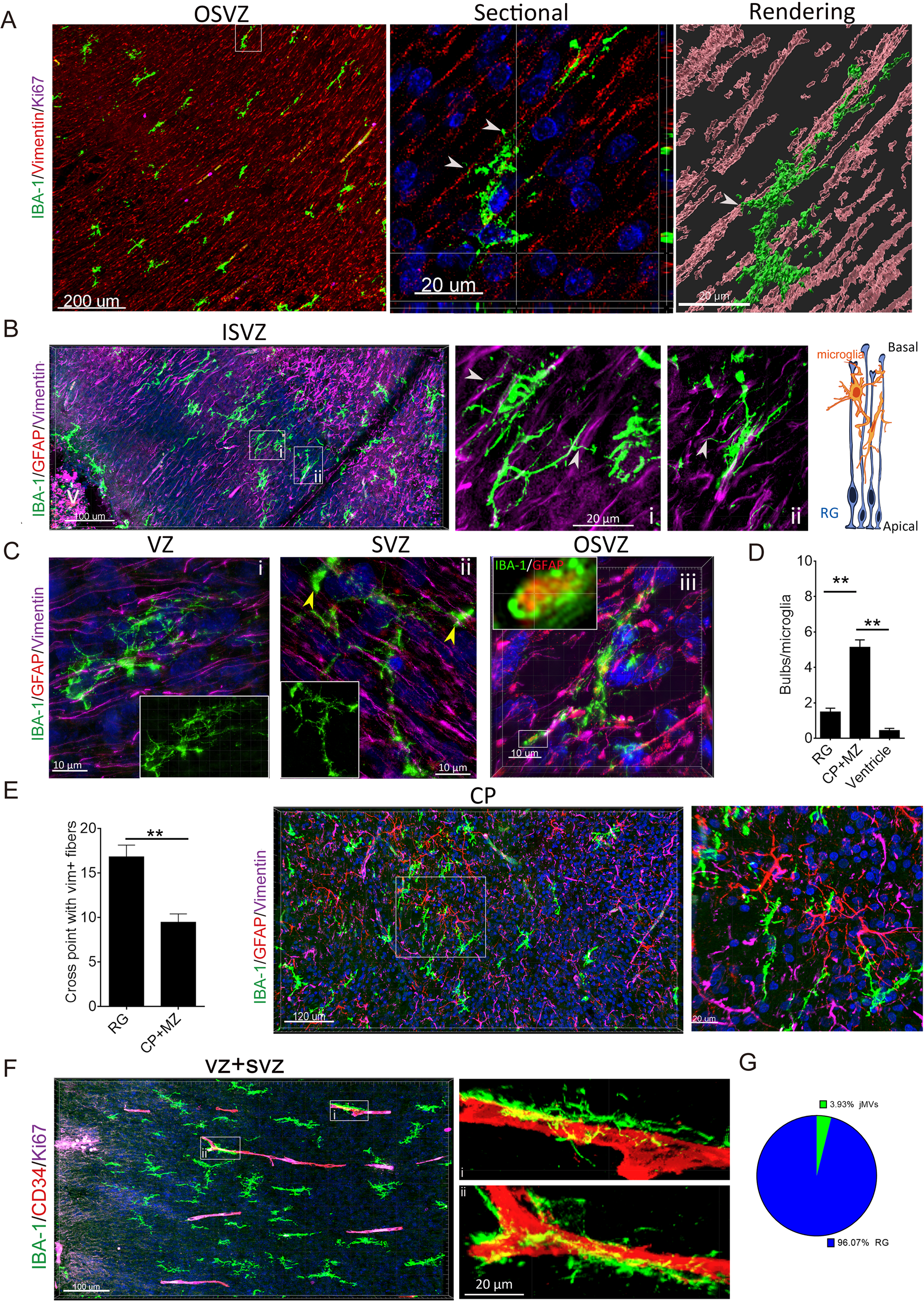
Human fetal microglia monitor and be scaffolded by RG fibers. **A.** Immunostaining of 13 gw fetal brains with IBA-1, Ki67, and vimentin antibodies in OSVZ revealed that the main axis of microglia paralleled the orientation of vimentin^+^ RG fibers. Most of the RG scaffolding microglia are negative in Ki67 staining. The sectional view in the middle panel and 3D rendering of Z-stacked images in the right panel show the microglia processes conjugated/scaffolded by RG fibers (white arrows, contact of microglia projections with vimentin^+^ fibers). **B.** Immunostaining of fetal brain with IBA-1, GFAP, and vimentin antibodies in ISVZ and VZ shows that the microglia nested between the RG fibers and their processes intensively contacted or crossed with the vimentin^+^ RG fibers (middle and right panel, magnified part of the boxed region; white arrows, the contact of the microglia projections with the vimentin^+^ fibers). The right panel is a graphical illustration of the relationship between microglia and RG fibers. V, ventricle. **C.** 3D SIM images of immunostaining with IBA-1, GFAP, and vimentin antibodies in VZ (i), SVZ (ii), and CP (iii) (yellow arrows, bulbous endings). The inner inserts in i/ii are IBA-1 immunostaining, showing the structure of the microglia. The inner insert in iii is the magnified image of the boxed region showing the GFAP^+^ engulfed fragment. **D.** Counting bulbous endings in RG scaffolding, CP, and ventricle microglia. Fetus: n = 4; mean ± s.e.m; *t-*test, **<0.01. **E.** Counting the microglial processes crossed with vimentin^+^ RG fibers in SVZ+VZ (RG), CP+MZ. Fetus, n = 4; Data, mean ± s.e.m; **; p < 0.01. The two panels on the right are representative images of IBA-1, vimentin, and GFAP antibodies immunostaining in the CP+MZ region of the 13-gw fetal brain. Microglia processes randomly cross with vimentin^+^ fibers, and the main axis of microglia lacks a specific orientation. **F.** IBA-1, CD34, and Ki67 antibodies immunostaining in VZ and SVZ of 13-gw fetal brain. Some microglia are scaffolded by the microvessels; most are negative in Ki67 staining. **G.** The ratio of RG scaffolding microglia and microvessel scaffolding microglia in the human fetal brain. Fetus, n = 4. Total images, n = 15

Microglia in the ventricular zone (VZ) and subventricular zone (SVZ), including inner SVZ (ISVZ) and OSVZ, nested between the radial glia (RG) fibers, and the main body axis of the microglia paralleled the orientation of the RG fibers **(Fig. 1A)**; the numerous processes of microglia were scaffolded by/ attached the RG fibers in 3D images **(Fig. 1A, B; right panels and rendering image)**. Most RG scaffolding microglia were non-proliferative **(Fig. 1A)**. 3D SIM images with 64-nm resolution revealed that the RG scaffolding microglia had several bulbous endings **(Fig. 1C-i, ii)**, which are structures for phagocytosis^3, 31^. Notably, we observed that some large microglia (>80 µm in the long axis) with multiple bulbous endings spanned a dozen RG fibers **(Fig. 1C-ii, iii)**. Moreover, the bulbous endings of the microglia contained GFAP^+^ debris **(Fig. 1C-iii, inner panel)**; the microglia in the cortical plate (CP) showed more bulbous endings than the RG scaffolding microglia **(Fig. 1D)**; the processes of RG scaffolding microglia had more crossing-points with the vimentin^+^ processes than those of the microglia in the CP regions **(Fig. 1E)**, reflecting that the migrating microglia were scaffolded by the RG fibers. However, only 4% of the microglia migrated to ISVZ and OSVZ were scaffolded by microvessels **(Fig. 1F, G)**. These findings indicate that migrating microglia with limited proliferative ability are primarily scaffolded by RG fibers and monitor them in the early fetal brain.

### Highly proliferative microglia aggregates with unique morphological features exist in the fetal brain

While screening the immunostained 50 µm sections, we identified a larger and denser aggregate of microglia in the CTIP2^+^ region close to the basal caudate in the fetal brain over 13 gw **(Fig. 2A-C; Fig. S2A)** and the density of microglia was far higher than that of the microglial clusters that others observed^16, 30^. Mouse whole-brain staining revealed that similar microglial aggregates were not present in the mouse fetal brain at 12.5 days **(Fig. S2B, C)**. The timeframe of the appearance of microglial aggregate coincides with the appearance of OSVZ in human fetal brains^13^. Therefore, we hypothesized that the aggregate might be a secondary microglial formation center with a robust proliferative ability that compensates for the lack of microglia during the rapid expansion of the human fetal brain after the formation of OSVZ. Ki67^+^ microglia in the aggregates accounted for 23.5% (2592/11034; n = 6) of the total microglia and lacked a specific orientation, while scattered Ki67^+^ microglia in other regions, including VZ, SVZ, and CP, accounted for 6.24% of total microglia **(Fig. 2D; Table S1)**. The ratio of Ki67^+^ microglia in the aggregates did not depend on gw **(Fig. 2E)**.

**Fig. 2.**
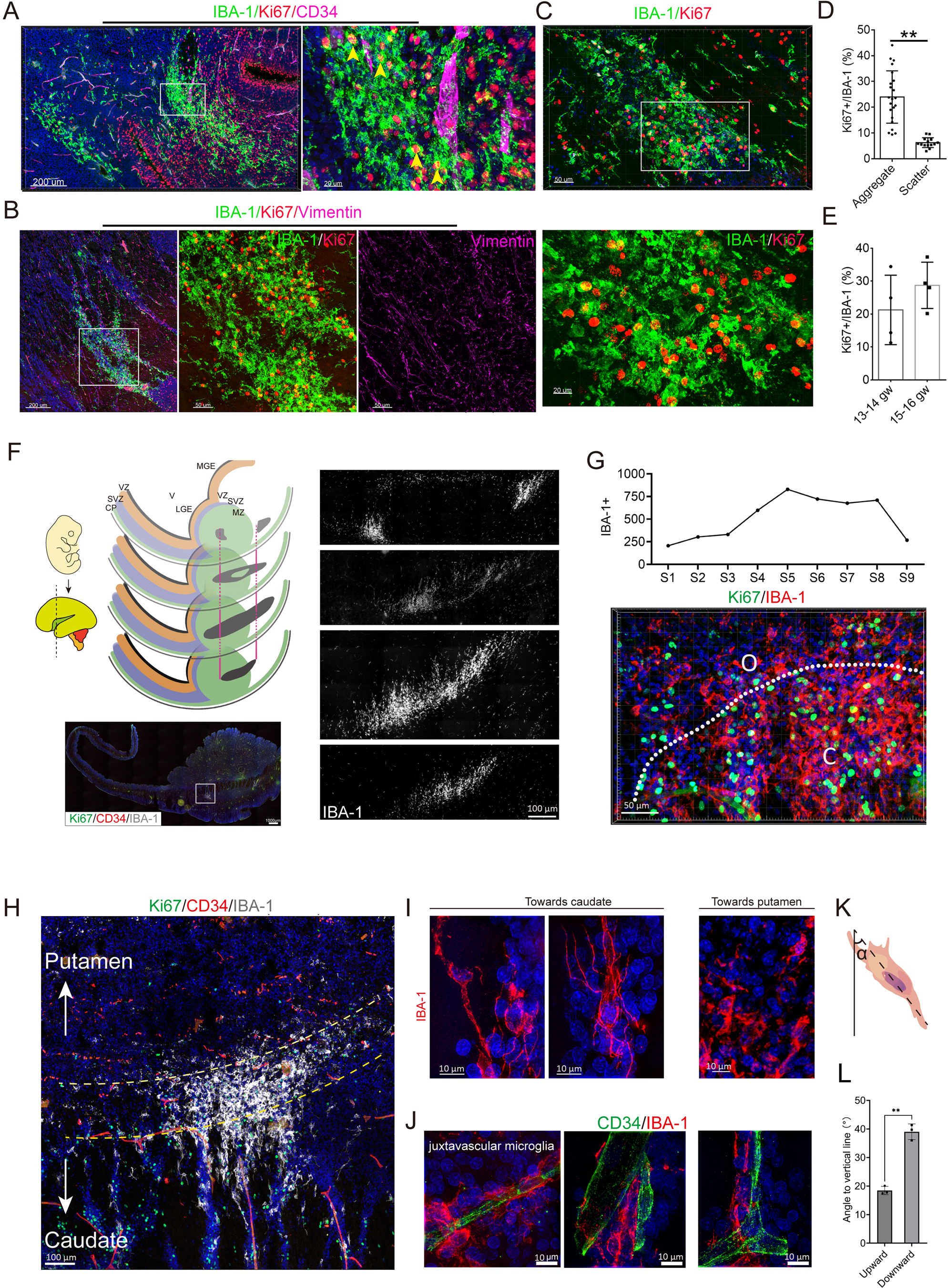
Identification of proliferative microglia aggregates in the fetal brain. **A-C**. 3D stitching images of IBA-1, Ki67, and vimentin (A) or CD34 (B) antibodies immunostaining in two fetal brains (13−14 gw) revealed a large microglia aggregate with abundant Ki67^+^ microglia. The orientation of vimentin^+^ fibers is chaotic in the center of the aggregates (right panel) (B). 3D stitching images of IBA-1 and Ki67 antibodies immunostaining in 13 gw fetal brains reveal an array of Ki67^+^ microglia aggregates (C). **D.** Percentage of Ki67^+^ microglia in aggregate and scattered microglia (scattered microglia in other regions). Fetus, n = 6; Data, mean ± SD **E.** Percentage of Ki67^+^ microglia in human fetal brain samples with 13−14 gw and 15−16 gw. Fetus, n = 6; Data, mean ± SD; *t-*test. **F.** The illustrated graph on the left indicates the anatomical position of the sections to estimate the size of the proliferative aggregates. The lower panel showed an image of a stitched whole section. LGE, lateral ganglionic eminence; MGE, medial ganglionic eminence; V, ventricle; CP, cortical plate; VZ, ventricular zone; SVZ, subventricular zone; MZ, mantle zone. The right panel shows the characteristics of the aggregate. **G.** Counts of IBA-1^+^ microglia in nine consecutive sections of the fetal brain 15 gw. The lower panel shows 3D high-resolution stitched images of Ki67/IBA-1 antibodies immunostaining, which revealed that the center harbors more Ki67^+^ microglia. The dashed line is the margin of the center of the aggregate and the outer region (O, outer; C, center). **H.** The shapes of microglia aggregate. Arrows are the orientation. The yellow dashed line is the main distribution area of IBA-1^+^ cells. **I, J.** Morphology of microglia in the border regions of proliferative microglial aggregates. The microglia on the border facing the putamen (superficial) lack long-oriented processes; the microglia on the border facing the caudate (deeper) show orientation and multiple extended processes (I). Microvessel scaffolding microglia at the border of the caudate (J). **K, L.** Graphical illustration of the angle between the vertical line representing a straight line from the meninges to the ventricle and the cellular long-body axis (K). The average angle between the body and vertical axes was 17−20° on the caudate surface; the average angle ranged 35−41° on the putamen-facing side (L). The vertical line is the line from the meninges to the ventricles. Fetus, n = 3. All data, mean ± S.E.M; **, p < 0.01.

By estimating the size of microglial aggregates in nine consecutive sections of 15 gw fetal brains, we found that the total estimated size reached 1.736 mm^2^ **(Fig. 2F, G)**, indicating that the microglial aggregate covered a larger area in the fetal brain. We uncovered a specific pattern of microglial aggregates: the microglial density gradually decreased from the center to the two ends **(Fig. 2G, upper panel)**. The center of microglial aggregates harbored more Ki67^+^ microglia with amoeboid morphology **(Fig. 2G, lower panel)**. However, the microglia were ramified or bipolar in the outer region of the aggregates **(Fig. 2I)**. Extended bipolar morphology is a signature of migrating microglia^32^, and a gradient of chemokines (e.g., calcium flicker) defines the orientation of microglia by changing the angle of the long axis^33^. In addition, the microglia in the outer region of the aggregate were toward the deep brain region rather than the superficial layers **(Fig. 2H)**, and were robustly scaffolded by microvessels on the surface of the caudate **(Fig. 2J)**. The long axis and acute angle measurements indicated that the microglia in the outer region faced the deep caudate (20°) rather than the superficial MZ (40°) **(Fig. 2K, L)**, suggesting some degree of directional alignment of the microglia in the aggregates ^34^ ^35^. Microglia, which are scaffolded by microvessels in the outer region of aggregates, also exhibit a distinct orientation toward the deep caudate rather than the superficial regions.

3D high-resolution images with stitching revealed that the short processes of amoeboid microglia in the aggregates rarely crossed with vimentin^+^ RG fibers, which are common in RG scaffolding microglia and did not exhibit long and complex processes in the CP region or adult human cortex **(Fig.3A-C)**. 3D ultra-high-resolution images with 64 nm resolution further showed that the microglia in the center were uniformly round and lacked bulbous endings and stem bulbs in the long projections that are common in RG scaffolding microglia and CP microglia, as well as membrane ruffles in ventricle microglia or macrophages exposed to growth factors and bacterial products **(Fig. 3D)**^36,37^. Strikingly, Ki67^+^ RG scaffolding microglia or ventricle microglia had extended/branched processes and bulbous endings or multiple processes and membrane ruffles compared to Ki67^+^ amoeboid microglia in microglial aggregates. Together, the microglia in the aggregates had distinct characteristics of proliferative ability and morphological and orientation identities.

**Fig. 3.**
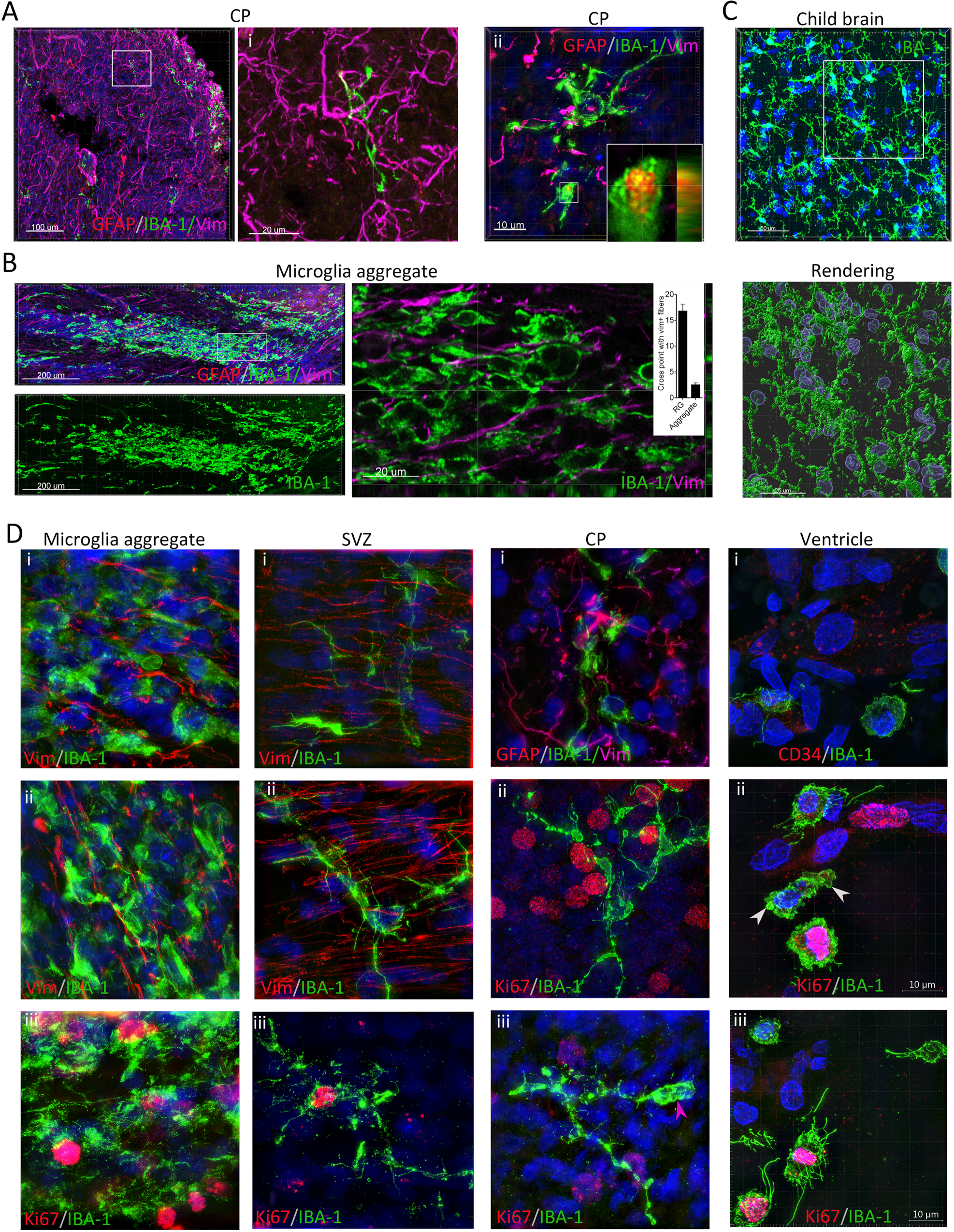
Microglia in proliferative microglia aggregates have a distinct morphological signature. **A-C.** High-resolution 3D stitching images of IBA-1, vimentin antibodies immunostaining revealed the morphology of microglia in CP (A), microglia aggregates (B), and child brain (C) and their relationships with vimentin^+^ RG fibers (63x oil objective). The microglia processes randomly cross with vimentin^+^ fibers in CP (inner bar graph); the bulbous ending of the microglia in CP contains GFAP^+^ debris (inner insert in ii, debris). Round and amoeboid microglia nested in vimentin^+^ fibers (right panel, sectional view). The Z-stacked image of the microglia in the cortex of the child’s brain reveals that the microglia processes form a large and complicated web-like structure (C). **D.** SIM images of IBA-1, vimentin or CD34, and GFAP or Ki67 antibodies immunostaining in proliferative microglia aggregates, SVZ, CP, revealed the distinct morphology of microglia in microglia aggregates. The microglia in the proliferative aggregate are uniformly round or amoeboid, including proliferative or non-proliferative; the RG scaffolding microglia have long projections spanning a dozen of RG fibers (i, ii); RG scaffolding Ki67^+^ microglia along exhibit long projections and bulbous endings (iii); the microglia in ventricle have membrane ruffles and Ki67^+^ microglia in the ventricle exhibit several slender processes and membrane ruffles (white arrows, membrane ruffles; pink arrows, bulbous ending).

### Introducing microglia into brain organoids replicates the proliferative microglia aggregate and enhances organoid complexity

YS-derived microglia enter the fetal brain at 4.5−5.5 gw^6^. After generating hiPS-derived cortical organoids (hCO), we introduced SV40 microglia with stable GFP expression into hCO on day 30, which were structurally and efficiently cross-talked with 2D neurons in the co-culture **(Fig. S3)**. The microglia were introduced into the hCOs by suspending them in a basal culture solution containing SV40 microglia **(Fig. 4A)**. While maintaining hCOs in the medium with SV40 microglia, we noticed that SV40 microglia were recruited into the hCOs **(Fig. 4A)** and formed a decent cluster on the hCO surface on day 7 **(Fig. 4B)**. Similar microglial aggregates appeared on the surface of the chimeric hCO with induced microglia^27^. During monitoring, we noticed that the morphology of microglia changed after attachment to the surface of hCO **(Fig. 4A and Fig. S4A)**. The microglia in the aggregates on the surface of the hCOs were highly proliferative and 38.2% (100/277) of the microglia in the aggregates expressed Ki67 **(Fig. 4B)**. However, hiPS-derived macrophages were not sufficiently recruited to the surface of hCO and did not form an identifiable cluster on the surface of hCO using the same protocol **(Fig. 4B-D)**. Histology revealed that bipolar microglia, but not induced macrophages, penetrated the hCOs sufficiently and deeply and were present at the edge of the VZ and neural tube **(Fig. 4A, B; see also other images)**.

**Fig. 4.**
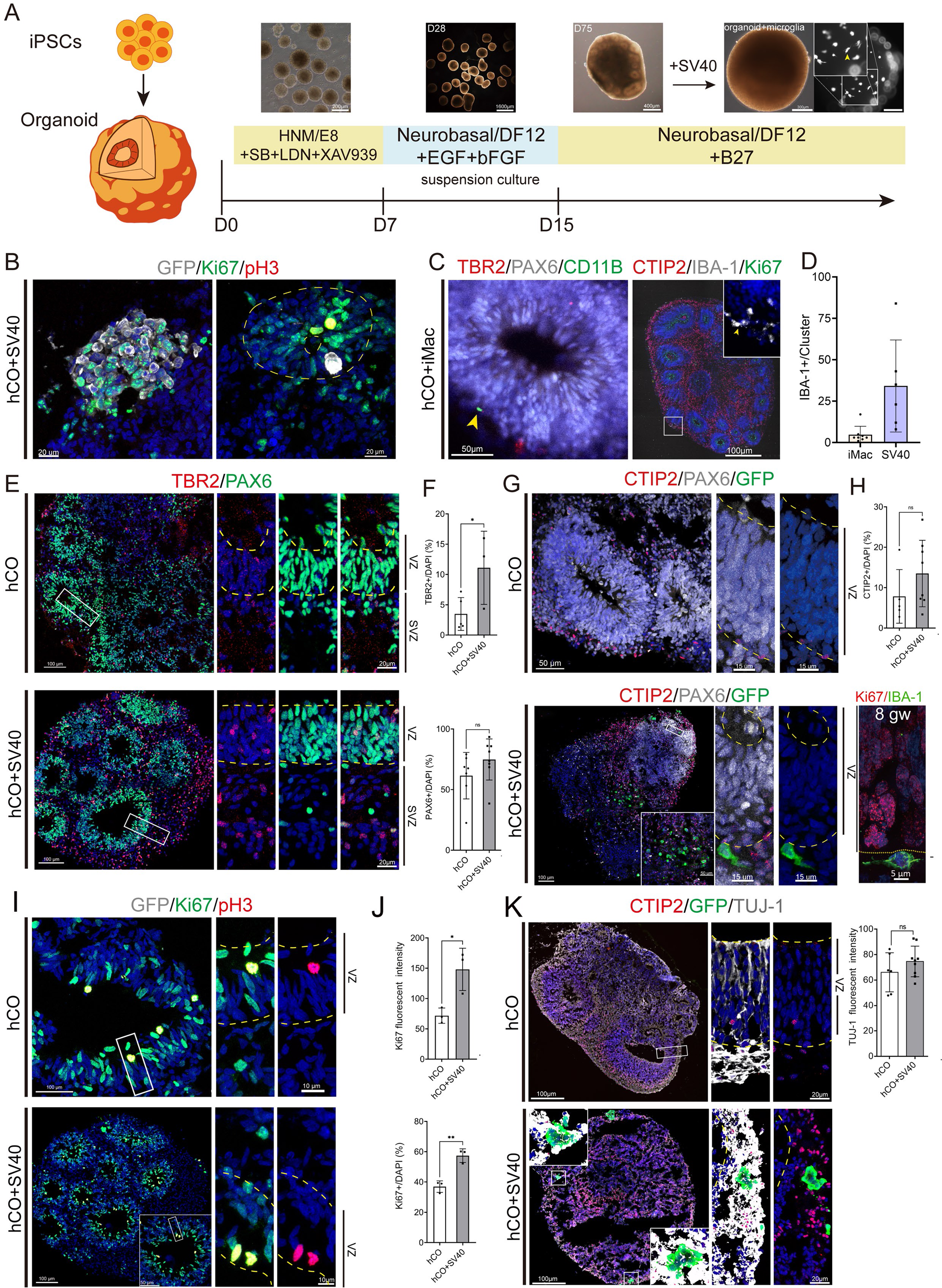
Introducing SV40 microglia into hCO replicates the proliferative microglia aggregate and enhanced neuronal maturation. **A.** The method of introducing SV40 microglia into hiPSCs-derived hCO. Microglia are introduced on day 30 by suspending the medium with SV40 microglia. Epifluorescent microscopy showed that the microglia are recruited to the surface of the hCO and that the morphology of the microglia is changing. **B.** Immunostaining with pH3, Ki67, and GFP antibodies in chimeric hCO with SV40 microglia on day 37 shows that Ki67^+^ microglia are enriched on the near inner surface of hCO and form an aggregate. Some of the SV40 microglia are present in the VZ region. **C.** Immunostaining with TBR2, PAX6, CD11B or CTIP2, IBA-1, Ki67 antibodies in chimeric hCO with iMac (hiPSCs-derived macrophages) co-cultured for 7 days. iMAC was added on day 30. A limited number of iMacs were recruited to the hCO but did not form aggregates on the surface of the hCO. **D.** Comparison of the count of IBA-1^+^ SV40 microglia on the surface of hCO with iMAC. Organoid, n ≥ 3. All data, means ± S.E.M; *, p < 0.05. **E.** Immunostaining chimeric hCO with SV40 microglia or hCO by PAX6, TBR2 antibodies revealed that more TBR2^+^ cells are shown in the SVZ region of hCO with SV40 microglia. The boxed region in the left panel is the magnified region in the right panel. **F.** Comparing the ratio (%) of TBR2^+^ or PAX2^+^ cells in hCO with SV40 microglia with hCO. Organoid, n ≥ 3. All data, means ± S.E.M; *, p < 0.05. **G.** Immunostaining images in hCO with SV40 microglia or hCO with CTIP2, GFP, and PAX6 antibodies. **H.** Comparison of the ratio (%) of TBR^+^ cells in hCO with SV40 microglia with hCO. Data, mean ± S.E.M; *t-*test. Sample size: n = 16 **I.** Immunostaining with Ki67, pH3, and GFP antibodies in hCO with SV40 microglia or hCO. **J.** Comparison of the ratio (%) or Ki67 intensity of Ki67^+^ cells in hCO with SV40 microglia with hCO. **K.** Immunostaining with CTIP2, GFP, and Tuj-1 antibodies in hCO with SV40 microglia introduced microglia on day 60 and hCO (yellow arrows, GFP^+^ SV40 cells).

The introduction of microglia into brain organoids promotes neuronal maturation and axon development^27^. Immunostaining revealed that the introduction of microglia into hCO on day 30 accelerated the formation of TBR2^+^ cells without significantly affecting PAX6^+^ progenitors in the VZ or CTIP2^+^ cells in the SVZ **(Fig. 4E-H)**. Furthermore, the introduction of microglia into hCO also increased the number of Ki67^+^ progenitor cells in neural tubes **(Fig. 4I-J)**. The chimeric hCO with microglia showed a denser and larger Tuj-1^+^ area than the hCO **(Fig. 4K). The introduction of** microglia into hCO on day 60 reduced PAX6^+^ progenitors but did not significantly affect CTIP2^+^ and Ki67^+^ cells in hCO **(Fig. S4B-D)**. In particular, amoeboid/bipolar SV40 microglia were present in the VZ region of the hCOs, similar to fetal microglia in the VZ **(Fig. 4G; lower left; Fig. S4E)**. These findings are consistent with the morphology of microglia derived from erythromyeloid progenitors (EMPs) or iPS in brain organoids in vitro^27, 38, 39^.

### Introducing microglia into hCO enhanced immunity and neuronal maturation

Necrosis in the brain activates microglia^40^. Brain organoids often have necrotic regions in the center^41^. Microglia are immune cells that secrete multiple immune-related cytokines that modulate brain immunity in the brain^42^. Nuclear fragments or MAP2^+^ fragments/PSD95^+^ puncta were present in the penetrating microglia of hCO, suggesting that the penetrating microglia of the hCO were phagocytic **(Fig. 5A; Fig. S3G)**. However, the penetrating microglia covered a smaller region but showed a larger volume **(Fig. 5B, C)**. Immunologically, the introduction of SV40 microglia into cortical organoids increased IL-1β, IL-8, CCL2, and CX3CL1 secretion and decreased TGF-β secretion **(Fig. 5D)**. Stimulation of hCO containing SV40 microglia with INF-γ, TNF-α, and LPS slightly increased IL-1β, IL-8, CCL2 and CX3CL1 secretion and slightly decreased TGF-β secretion compared to non-treated organoids. However, these changes were not significant **(Fig. 5D)**. Collectively, the microglia in the hCOs were phagocytic and inflammatory.

**Fig. 5.**
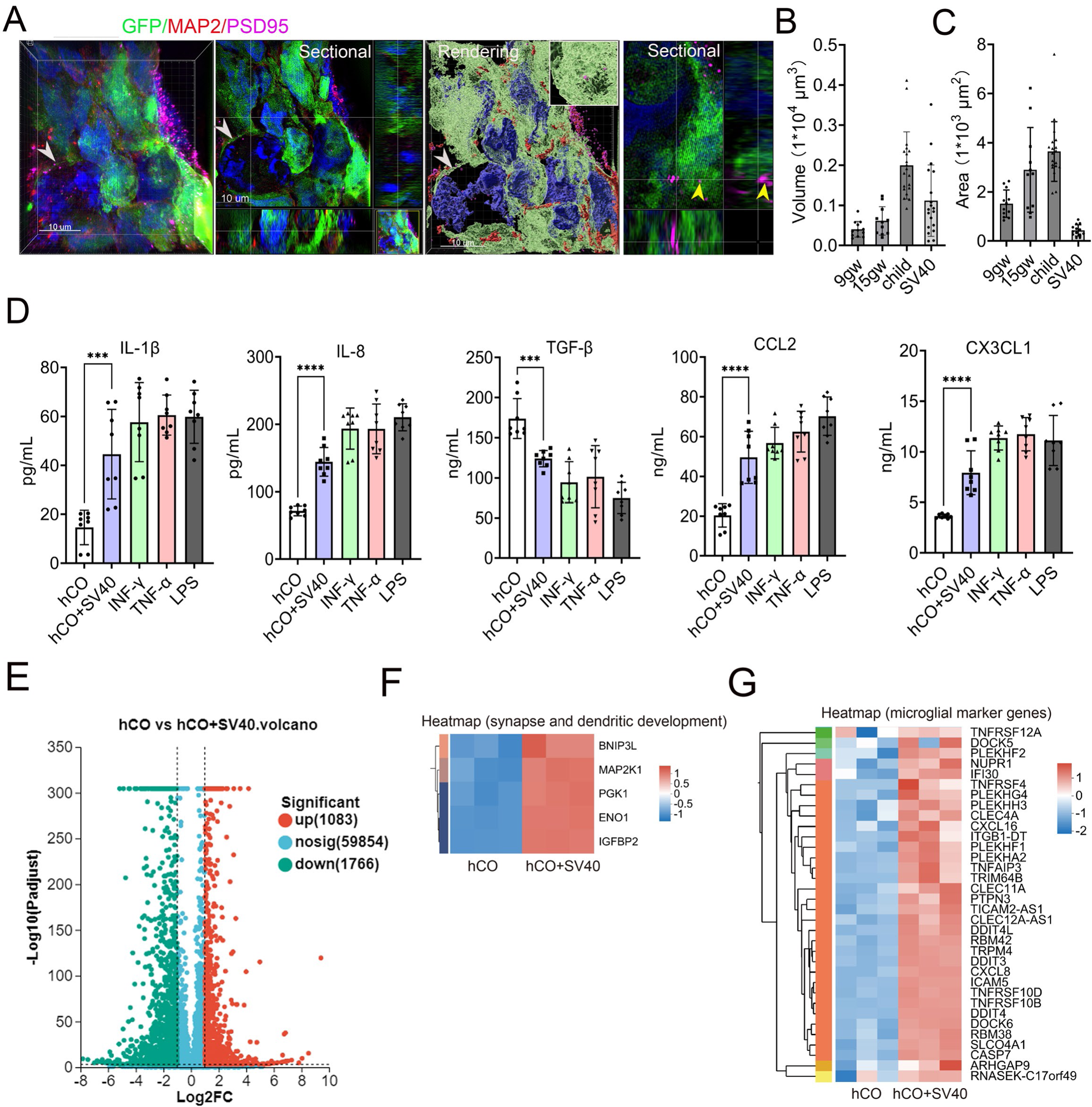
The introduction of SV40 microglia into hCO increased immunity and neuronal maturation. **A.** Immunostaining of hCO with SV40 microglia cultured for 120 with IBA-1, PSD95, and MAP2 antibodies. The two left panels revealed fragmented nuclear debris shown in the penetrated SV40 microglia (white arrows, fragmented nuclear debris; yellow arrows, the right two panels show that PSD95^+^ puncta are included in the SV40 microglia. **B, C.** Comparison of the volume and size of penetrated microglia with the microglia in the fetal brain with 9.5 and 15 gw and a child brain aged 6 years. SIM images are used for volume and size analyses. Data, mean ± S.E.M; *One-way ANOVA*. **D**. IL-1ß, IL-8, TGF-ß, CCL2, CX3CL1 levels in the supernatant of chimeric hCO with SV40 microglia and hCO or those after treating with IFN-γ, LPS and TNF-ɑ. n ≥ 4. Data, mean ± S.E.M; *One-way ANOVA*; *, p < 0.05, **, p < 0.01; ns, no significant difference. **E** Volcano plots of upregulated or downregulated genes in hCO with SV40 microglia. **G, H.** Heatmaps show genes that are upregulated in chimeric hCOs encoding microglial-related cytokines and proteins.

To gain deeper insight into the roles of microglia in organoids, we performed RNA-seq analysis for hCO after introducing microglia for 7 days and identified changes in 2849 genes, including 1083 upregulated genes and 1766 downregulated genes **(Fig. 5E)**. We found that a group of genes related to synaptic and dendrite developments^43^, including *BNIP3L, MAP2K1, PGK1, ENO1,* and *IGFBP2* genes, were upregulated in hCO with SV40 microglia **(Fig. 5F)**. In addition to microglia-specific marker genes, *CXCL16*, an anti-inflammatory cytokine in microglia^44^, was upregulated in hCO with SV40 microglia **(Fig. 5G)**. Consistent with the results of the supernatant analysis, CXCL8 was upregulated in hCO with SV40 microglia **(Fig. 5G)**. We identified microglial marker genes, including *NUPR1, TNFAIP3, TNFRSF19, CLEC11A, and CLEC4A*, in cortical organoids with microglia. *NUPR1*, a receptor for oncostatin, increases microglia-specific cytokine levels during injury ^45^. *TNFAIP3 and TNFRSF19* regulate processes such as apoptosis and growth and have anti-inflammatory effects on microglia^46, 47^. *CLEC11A and CLEC4A* encode immune response-associated proteins or chemokines^48–50^.

### DS Fetal brain exhibited increased microglia phagocytic structures and expanded proliferative microglia aggregates

To determine the status of proliferative microglial aggregates in the fetal brain with impaired neuronal development, we analyzed a 16 gw DS fetal brain, characterized by altered cortical development and impaired RG^51^, using IBA-1, CD34 and Ki67 immunostaining. We identified a larger microglial aggregate in DS fetal brain compared to healthy fetal brains (DS vs. healthy: 4.168 vs. 0.108−2.129 mm^2^), and the morphometry and percentage of Ki67^+^ microglia in the aggregates were nearly identical to those of healthy fetal brains **(Fig. 6A-E; Fig. S5A)**. The microglia in the aggregates of DS fetal brain were morphologically identical to those of healthy fetal brain in the SIM images and lacked long processes, bulbs, and membrane ruffles **(Fig. 6B)**. 3D ultrahigh-resolution images revealed that most microglia in brains with DS, including RG scaffolding, CP, and microvascular scaffolding microglia, had more complicated processes, bulbous endings, and stem bulbs (**Fig. 6F-H; Fig. S5B-D)**. The bulbous endings of DS microglia had higher levels of clathrin^+^ puncta than those of the processes of microglia without bulbs **(Fig. 6I)**. Consistent with the high microglial phagocytic signatures in DS fetal brain, more caspase 3^+^ distorted neuronal projections and neuronal bodies with fragmented nuclei were present in DS fetal brain **(Fig. S6A, B)**. Expanded microglial aggregates in DS and healthy fetal brains did not harbor caspase 3^+^ cellular processes and cellular bodies, CD8^+^ cells, or CD177^+^ activated neutrophils **(Fig. S6A, C)**. These findings indicate that a larger microglial aggregate is related to impaired neuronal development in the DS fetal brain and differs from an inflammatory aggregate of immune cells.

**Fig. 6.**
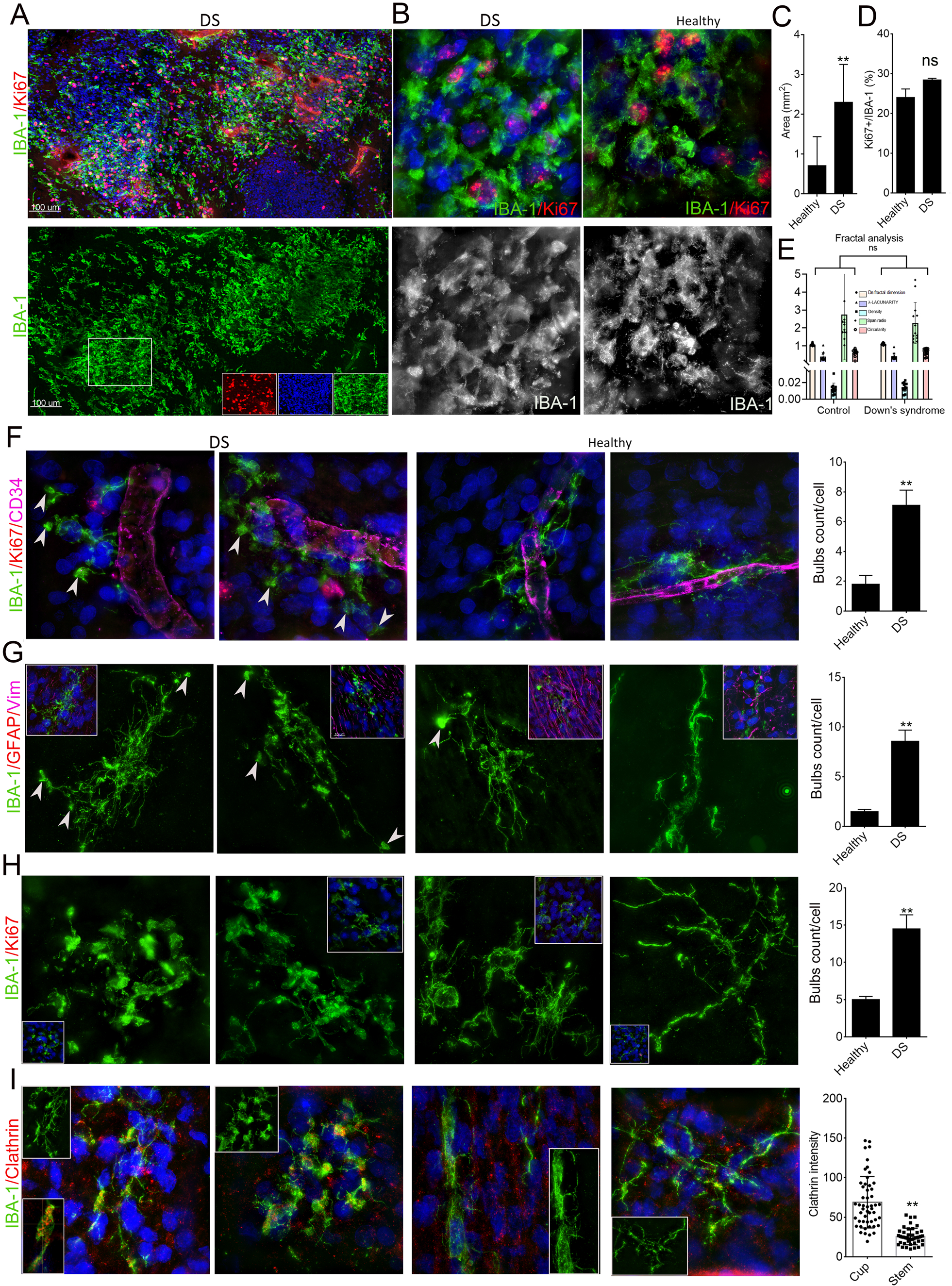
Proliferative microglia aggregates expanded in the DS fetal brain with increased neuronal death and intensified microglia phagocytosis. **A.** Representative images of IBA-1, Ki67, and CD34 antibodies immunostaining in DS fetal brain with 16 gw reveal larger proliferative microglia aggregates. **B.** SIM images of IBA-1, Ki67, and CD34 antibodies immunostaining show that the morphology of the microglia in the center of the microglia aggregate is amoeboid and identical to the microglia of the aggregates in a healthy fetal brain. **C.** Comparison of the size of microglia aggregates in the DS fetal brain with those of the healthy brain. Data, mean ± S.E.M. t-test, **, <0.001. **D.** Comparison of the Ki67^+^ microglia of microglia aggregates in the DS fetal brain with that of the healthy brain. Data, mean ± S.E.M. t-test, ns, non-significant. **E.** FracLac analysis of microglia in the microglia aggregates of DS fetal brain and healthy brain revealed that the morphometry of microglia in the aggregate of DS fetal brain is nearly identical to that of healthy fetal brain. Data, mean ± s.e.m. t-test, ns, non-significant. **F.** SIM images of IBA-1, Ki67, and CD34 antibodies immunostaining revealed that microvessel scaffolding microglia in DS fetal brain have more bulbous endings (white arrows, bulbous endings). The bar plot is the count of bulbs for each microglia. Data, mean ± S.E.M. *t*-test, **, <0.001. **G.** SIM images of IBA-1, Ki67, and CD34 antibodies immunostaining revealed that RG scaffolding microglia in DS fetal brain have more bulbous endings (white arrows, bulbous endings). The bar plot is the count of bulbs for each microglia. Data, mean ± S.E.M. *t*-test, **, <0.001. **H.** SIM images of IBA-1, Ki67, and CD34 antibodies immunostaining revealed that CP microglia in DS fetal brain have more bulbous endings and stem bulbous structure. The bar plot is the count of bulbs for each microglia. Data, mean ± S.E.M. *t*-test, **, <0.001. **I.** SIM images of IBA-1 and clathrin antibodies immunostaining revealed that CP microglia in DS fetal brain have higher levels of clathrin puncta in the bulbous structure. Data, mean ± S.E.M. *t*-test, **, <0.001.

## Discussion

YS-originating microglia penetrate the neural tube in the human fetal brain before the appearance of OSVZ, thus contributing to a large human cortex. Despite rigorous studies in rodents that reveal microglialimmune pathophysiology and colonization in the developing and diseased brain^11^, human-specific microglial colonization in the fetal brain is unknown. The density of microglia in the brain is modest among different species, while human microglia maintain homeostasis in the largest, most complex, and most intellectual brains on earth^52^. Therefore, it is reasonable to speculate that the total number of microglia in the human brain is the highest among most species. Multiple clusters of microglia with amoeboid morphology that were only proliferative before 12 gw were observed in the human fetal brain, considering that these clusters, “a waiting site,”, are involved in eliminating lavish axons, driving the growth of axons, and directing the pathway finding of the axon^6, 30, 53^. The cluster resolves around 32 post-conceptual weeks^51^. The highly proliferative microglial aggregates found in this study could compensate for the lack of microglia during the rapid expansion of the fetal brain after the appearance of the OSVZ, especially in deeper brain regions. Microglia are phagocytes that differ from other macrophages in their origin and molecular identities^5^. The surface aggregate formation of microglia in chimeric hCO revealed that aggregation and proliferation were unique characteristics of microglia, but not of macrophages.

Morphology is a common standard to assess the microglial status of microglia^54^. The morphology of microglia in aggregates lacks long processes, membrane ruffles, and bulbs, suggesting that the microglia in aggregates differ from other proliferative microglia and might not be involved in the elimination lavish axons and apoptotic bodies, driving the growth of axons, or directing the pathway finding of the axon^6, 30, 53^. In addition, the aggregate did not include other immune cells, such as cytotoxic T cells and activated neutrophils, in DS and healthy brains, indicating that microglial aggregation differs from immune cell aggregation in inflammation, which is characterized as an aggregate of immune cell diversity. The orientation of the microglia to the deep brain region highlighted that the migrating microglia from the secondary microglia formation centers populated the deep brain region rather than the superficial layers of the fetal brain.

A limitation of this study is that we did not have more evidence on the extent to which highly proliferative aggregates contribute to the precise and full colonization of the human fetal brain. If the absence or insufficiency of proliferative microglia aggregates is related to human-specific brain behaviors or disorders such as higher cognitive ability or autism and schizophrenia et al.^55^, it is unknown. Human microglia maintain homeostasis in the brain and prune excess synapses^11^. The aggregate expanded in the DS fetal brain. Therefore, it is rational to envision that human-specific proliferative microglial aggregates could be associated with mental disorders related to brain development. A small number of ramified microglia or amoeboid microglia with membrane ruffles and bulbs were proliferative. Thus, proliferative microglia aggregates might partially compensate for the shortage of microglia, but not for the renewal of microglia. However, the cues that drive the formation of proliferative aggregates remain unknown. Microglia in 2D culture, including co-culture of microglia with neurons, did not form an obvious aggregate, suggesting that secondary microglial formation centers require the 3D structure of the brain.

We identified a secondary microglial formation center in the human fetal brain that expanded in the DS fetal brain with impaired neuronal development and paralleled the development of OSVZ. We also replicated the secondary microglial formation center in chimeric brain organoid models using microglia, suggesting that the formation of the secondary microglial formation center is a unique characteristic of human microglia and the human fetal brain.

## Data availability

All relevant data are within the paper and are available to anyone for a reasonable request.

## Author contributions

CY.S and H.S designed and performed most of the experiments, analyzed the data and wrote the paper.

XY.C and Y.L performed the construction of organoids.

QZ.H and L.G collected human fetal brain samples.

R.J tested microglia.

LX.M provided financial support and supervised the project.

