## Supplementary material for "Secondary microglia formation center in the human fetal brain": methods, extended figure legends/figures, supplemental table S1

### Extended Materials and Methods

#### hiPSCs/hESCs and human fetal brain samples

The ethics of human subjects were approved by the Ethics Committee of the Institutes of Biomedical Sciences at Fudan University (No. 28). Yamanaka's factors (Oct4, Sox2, Klf4, and c-Myc) were applied to construct hiPS cell line (8-12). Human ES cell line H9 (WA09) (provided by Prof. Su-Chun Zhang) and hiPS cell line (8–12) were cultured in a feeder-free culture and maintained in Matrigel-precoated (Corning; CAT#:354234) or Vitronectin XF-precoated (STEMCELL; CAT#:07180) 6-well plates for long-term. Dr. Qizhi collected fetal brain tissue in Shanghai First Maternity and Infant Hospital with informed consent forms signed by each patient (2023-209).

#### Neural differentiation in 2D

Based on our previous protocol[1], ES/iPS cells were dissociated into small clumps in appropriate sizes using dispase. After a seven-day suspension culture, the embryoid bodies (EB) were subsequently transferred to an adherent culture dish with 10% serum for incubating for another 12 hours. On day 15, the part of the adherent rosette in the center was gently separated and collected for further differentiation.

On day 30, the cells were dissociated into a single unit using Accutase (Gibco, CAT#: A1110501), re-suspended, and evenly plated on the slides for further adherent culture. During the first 25 days, the basal culture medium comprised 96% DMEM/F12 supplemented with 1% Non-Essential Amino Acids (NEAA), 1% Glutamax, 1% N2 supplement, and 1% Penicillin-Streptomycin (PS). The following agents were added in the first 7 days: 2 μM SB-431542 (Ametek Scientific, CAT#: DM-0970), 0.3mM LDN-193189 (STEMGENT, CAT#:040074). From Day 10 to 25, the culture medium was continuously supplemented with 0.65 μM purmorphamine (PUR), a ventral morphologene[2]. After Day 25, 96% Neurobasal medium supplied with 1% NEAA, 1% Glutamax, 1% N2 supplement, 1% PS, and 2% B27 supplement is the basal medium for further maintenance.

**Induction of iPSCs to induced macrophages**

The differentiation method refers to the previously reported protocol with minor adjustments[3]. Briefly, ESCs/iPSCs were treated with Versene (15040-066, Gibco) and transferred to low-attachment plates (3471, Corning) to form embryoid bodies (EBs) in mTeSR medium (85850, STEMCELL Technologies). EBs were cultured in BMP-4 (10 ng/mL) and bFGF (5 ng/mL) in APEL II medium (05270, STEMCELL Technologies) on day 1 to generate primitive streak-like mesodermal progenitor cells. From day 2 to 7, BMP-4 (10 ng/ml), bFGF (5 ng/ml), VEGF (50 ng/ml), and SCF (100 ng/mL) were supplied to achieve hematopoietic specification. On day 8 to 9, bFGF (10 ng/ml), VEGF (50 ng/ml), SCF (50 ng/mL), IGF-1 (10 ng/mL), IL-3 (25 ng/mL), M-CSF (50 ng/mL) and GM-CSF (50 ng/mL) were supplied to get myeloid lineage differentiation. At 10$^{th}$ day, the cells were transferred into StemProTM-34 SFM medium (10639011, Gibco) with bFGF (5 ng/ml), VEGF (50 ng/ml), SCF (50 ng/mL), IGF-1 (10 ng/mL), IL-3 (25 ng/mL), M-CSF (50 ng/mL) and GM-CSF (50 ng/mL) for culturing for another 10 days. From day 20 to 22, the floating cells were collected and plated on a Matrigel plate in myeloid maturation medium with bFGF (5 ng/ml), VEGF (50 ng/ml), SCF (50 ng/mL), IGF-1 (10 ng/mL), IL-3 (25 ng/mL), M-CSF (100 ng/mL) and GM-CSF (100 ng/mL). From day 22 to 27, IL-3 was withdrawn from the medium. From Day 28, iPSC-macrophages (iMac) are maintained in RPMI1640 medium with 10% FBS or H3000 medium containing M-CSF (100 ng/mL) and GM-CSF (100 ng/mL). All recombinant agents were purchased from PeproTech.

**hCOs culture**

hCOs were generated using our previous protocol[2]. The iPSC/ES lines were initially dissociated into single cells and transferred to a 96-well plate featuring an ultra-low adsorptive V-bottom. Morphogens were added at different time intervals to induce differentiation. The day of initial iPSC digestion into spherical structures is marked as Day 0. On Day 1, the spheroid hCO was retrieved from the 96-well plate and transferred into a suspending culture. From day 1 to 6, the culture system included the following components: 48% DMEM/F12, 48% E8 culture solution, 1% NEAA, 1% Glutamax, 1% N2 supplement, 1% PS, 2 μM SB-431542, 0.3mM LDN-193189. Optionally, the WNT pathway inhibitor XAV-939 (2.5 μM) and the two SMAD pathway inhibitors were supplied. From day 7 to Day 14, the culture medium included: 96% Neurobasal culture solution, 1% NEAA, 1% Glutamax, 1% N2 supplement, 1% PS, 20ng/ml EGF, and 20ng/ml bFGF. Since Day 15, the small molecules were withdrawn, and B27 was introduced to the base culture solution with 2% concentration.

**Human immortalized microglia (HMC3/SV40) culture**

Human immortalized microglia were purchased from Applied Biological Materials Inc. (www.abmGood.com, Cat. No. T3961) 1 ml of Applied Cell Extracellular Matrix (ACEM) was added to the walled six-well plate for passaging. Medium for SV40 is Prigrow III (Cat. TM003) or DMEM/F12 basal medium with 10% FBS serum. Passage or subsequent experiments (e.g., coculture) are performed when the cells reach about 80% in the six-well plate. Add 1 ml of trypsin preheated to 37°C and leave to digest for 1 minute in the incubator. Digestion was stopped by adding 1 ml of cytogenic solution, and the cell clones were blown using a pipette gun. The cell suspension was collected into a 15 ml centrifuge tube and centrifuged using a centrifuge for 1.5 min at 1500 rpm.

**Generation of chimeric hCO with SV40 microglia**

On day 30 and day 60 of the hCO culture, SV40 microglia were added to the medium of hCO at a density of 10,000 cells/ml after digestion and resuspension. The hCO and SV40 microglia coculture were moved to a shaker overnight to ensure efficient contact. The next day, the medium was replaced by another medium without microglia, maintained for 7 days.

**Histology and Immunostaining**

Organoids and fetal brain samples were fixed with 4% paraformaldehyde (PFA) for 0.5 h and 24 h, washed in PBS several times, transferred to a 30% sucrose for dehydration, and embedded in OCT for cryo-sectioning. The organoids sections with 15 μm and fetal brain sections with 50 μm were washed in PBS, blocked, permeabilized in blocking buffer (0.3% Triton X-100, 10% normal donkey serum in PBS) for 1 h, and incubated with primary antibodies in a buffer (PBS with 0.3% Triton X-100 and 5% donkey serum) overnight at 4 °C. After washing in PBS for 2 hours, the slices were incubated with a secondary antibody solution that contained the corresponding fluorescence conjugated antibodies, DAPI solution, and 5% donkey serum for 1 hour. After washing several times, the sections were mounted on an aqueous mounting medium for microscopy.

Table S2. Primary antibody list

| antibody | host | Cat. | concentration |
|---|---|---|---|
| BDNF | Sheep | AB1513p | 1:500 |
| Caspase3 | Rabbit | #9669 | 1:1000 |
| CD11b | Mouse | ab133357 | 1:15000 |
| CD177 | Mouse | MEM166 | 1:1000 |
| CD34 | Mouse | ab8536 | 1:1000 |
| CD8 | Rabbit | ab4505 | 1:1000 |
| Clathrin | Rabbit | ab172958 | 1:500 |
| CTIP2 | Rat | ab18465 | 1:1000 |
| DARPP32 | Rabbit | AB1656 | 1:1000 |
| GFAP | Mouse | ab279290 | 1:1000 |
| GFAP | Rabbit | Z0334 | 1:1000 |
| GFP | Chicken | ab6556-25 | 1:1000 |
| GSH2 | Rabbit | 3388451 | 1:500 |
| IBA-1 | Goat | ab5076 | 1:1000 |
| Ki67 | Rabbit | LV1825852 | 1:1000 |
| MAP2 | Mouse | m1406 | 1:1000 |
| MAP2 | Rabbit | sc20172 | 1:5000 |
| MASH1 | Mouse | 556604 | 1:500 |
| MEIS1/2 | Goat | sc-10599 | 1:500 |
| OTX2 | Goat | AF1979 | 1:1000 |
| PAX6 | Rabbit | 901301 | 1:1000 |
| PH3 | Mouse | 9706 | 1:1000 |
| PSD95 | Rabbit | ab18258 | 1:1000 |
| SOX2 | Mouse | MAB2018 | 1:1000 |
| TBR2 | Sheep | AF6166 | 1:500 |
| TUJ1 | Mouse | T8660 | 1:1000 |
| TUJ1 | Rabbit | PRB-435P | 1:10000 |

Table S3. Secondary antibody list

| antibody | company | concentration |
|---|---|---|
| Alexa Fluro 488 Donkey anti-mouse IgG | Invitrogen | 1:1000 |
| Alexa Fluro 594 Donkey anti-mouse IgG | Invitrogen | 1:1000 |
| Alexa Fluro 594 Donkey anti-rabbit IgG | Invitrogen | 1:1000 |
| Alexa Fluro 488 Donkey anti-rabbit IgG | Invitrogen | 1:1000 |
| Alexa Fluro 594 Donkey anti-goat IgG | Invitrogen | 1:1000 |
| Alexa Fluro 488 Donkey anti-rat IgG | Invitrogen | 1:1000 |
| Cy5 AffiniPure Donkey Anti-Goat IgG (H+L) | Jackson | 1:500 |
| Cy5 AffiniPure Donkey Anti-mouse IgG (H+L) | Jackson | 1:500 |
| Cy5 AffiniPure Donkey Anti-rabbit IgG (H+L)<br>Phalloidin-633 | Jackson<br>Sigma | 1:500<br>1:200 |
| DAPI | Sigma | 1:1000 |

**High-resolution images and image processing**

Nikon Structured Illumination Microscopy (SIM) (Nikon, Japan) and Zeiss 880 or 710 confocal microscopy (Zeiss, Jena, Germany) were applied to scan all images. ImageJ software (Fiji, NIH, Bethesda, MD, USA) was used for cell counting, fluorescence intensity statistics, and morphological analysis. Imaris 9.8 (Bitplane AG, Zürich, Switzerland) was used for cell morphological analysis.

**Cytokine stimulation assays**

20 ng/ml human interferon-gamma (INF-γ) (AF-300-02) and 20 ng/ml tumor necrosis factor-alpha (TNF-α) (AF-300-01A), and 20 ng/ml lipopolysaccharide (LPS) (L4130-100MG) stimulated the hCOs or chimeric hCOs for 72 h. Subsequently, the supernatants were harvested and switched into a fresh culture medium for 48 h. Cell supernatants were collected daily to detect cytokine levels.

**RNA extraction**

Total RNA was extracted from the tissue using TRIzol Reagent according to the manufacturer's instructions. Then, RNA quality was determined by 5300 Bioanalyzer (Agilent) and quantified using the ND-2000 (NanoDrop Technologies). Only high-quality RNA sample (OD260/280=1.8~2.2, OD260/230≥2.0, RIN≥6.5, 28S:18S≥1.0, >1μg) was used to construct sequencing library.

**Library preparation and sequencing**

RNA purification, reverse transcription, library construction, and sequencing were performed at Shanghai Majorbio Bio-pharm Biotechnology Co., Ltd. (Shanghai, China) according to the manufacturer's instructions (Illumina, San Diego, CA). The RNA-seq transcriptome library was prepared following Illumina Stranded mRNA Prep, Ligation from Illumina (San Diego, CA) using 1μg of total RNA. Shortly, messenger RNA was first isolated according to the polyA selection method by oligo(dT) beads and then fragmented by fragmentation buffer. Secondly, double-stranded cDNA was synthesized using a SuperScript double-stranded cDNA synthesis kit (Invitrogen, CA) with random hexamer primers (Illumina). Then, the synthesized cDNA was subjected to end-repair, phosphorylation, and 'A' base addition according to Illumina's library construction protocol. Libraries were size selected for cDNA target fragments of 300 bp on 2% Low Range Ultra Agarose followed by PCR amplified using Phusion DNA polymerase (NEB) for 15 PCR cycles. After quantified by Qubit 4.0, the paired-end RNA-seq sequencing library was sequenced with the NovaSeq Xplus sequencer (2 × 150bp read length).

**Quality control and Read mapping**

The raw paired-end reads were trimmed, and quality was controlled by fastp[4] with default parameters. Then, clean reads were separately aligned to the reference genome with orientation mode using HISAT2[5] software. StringTie6 assembled the mapped reads of each sample in a reference-based approach.

**Differential expression analysis and Functional enrichment**

To identify DEGs (differential expression genes) between two different samples, the expression level of each transcript was calculated according to the transcripts per million reads (TPM) method. RSEM[6] was used to quantify gene abundances. Essentially, differential expression analysis was performed using the DESeq2[7] or DEGseq[8]. DEGs with |log2FC| ≧ 1 and FDR < 0.05(DESeq2 ) or FDR < 0.001(DEGseq) were considered to be significantly different expressed genes. In addition, functional-enrichment analysis, including GO and KEGG, was performed to identify which DEGs were significantly enriched in GO terms and metabolic pathways at Bonferroni-corrected P-value <0.05 compared with the whole-transcriptome background. GO functional enrichment and KEGG pathway analysis were carried out by Goatools and Python script, respectively. Reactome functional enrichment was carried out by Python scipy. All data were analyzed on the online platform of Majorbio Cloud Platform (https://cloud.majorbio.com/)[9].

**Alternative Splice Event Identification**

All the alternative splice events in our sample were identified by recently using the releases program rMATS[10]. Only the isoforms that were similar to the reference or comprised novel splice junctions were considered, and the splicing differences were detected as exon inclusion, exclusion, alternative 5′, 3′, and intron retention events.

**Tissue clearing**

Whole-head 3D images refer to a CUBIC method[11]. Briefly, after rinsing the fixed mice head in PBS three times, the whole head was incubated in CUBIC reagent-1 for 5–7 days on a 37°C shaker to become transparent, washed in 5 ml of PBS for 2 h, removed into new bottles for IBA-1 and Ki67 antibodies (PBS with 10% donkey serum and 1% Triton-X 100) incubation on a 37°C shaker for 3 days, washed five times (1 h/time) in 5 ml PBS, transferred to a bottle with for secondary antibody (PBS containing 5% donkey serum and 0.5% Triton-X 100), incubated 24 h in a 37°C shaker, washed five times in 5 ml PBS (1 h/wash), and re-cleared in reagent-2 before mounting. The following agents were used: Reagent-1 consisted of water (35%, weight/weight), urea (25%, w/w), Quadrol (25%, w/w), and Triton-X 100 (15%, w/w). Reagent-2 comprised of water (15%, w/w), urea (25%, w/w), sucrose (50%, w/w), and triethanolamine (10%, w/w). The ovaries were fixed in ice-cold PFA for 24 h and then transferred to sugar (30%, w/v) for 24 h.

**Statistical analysis and software**

For the selection of statistical regions, we chose to capture multiple images of different regions of embryonic brains and the vicinity of the neural tube of organoids. For cell number quantification, we utilized Imaris "spots" (6.5 μm) or Image-J (Fiji) "cell counter" for statistical analysis.

All clip angle analysis, area statistics, and fluorescence intensity analysis were performed using Image-J (Fiji). After separate statistics for different iPSC cell lines and subgroups, the average data was a single point. Each group contained ≥3 organoids, with ≥3 images taken of each organoid. Statistics were then performed using GraphPad, followed by a t-test or One-way ANOVA. We considered the data to be significantly different when $P \leq 0.05$.

**The structural analysis of microglia**

Structural analyses, including skeleton and FracLac analyses, were performed using Image-J (Fiji). Immunofluorescence staining images of a single were imported into the software, changed to 8-bit type images, and processed by "Process-Binary-make binary" for skeletonization. Skeleton analysis was subsequently performed using the "analyze-skeleton-analyze skeleton."

For FracLac analysis, the panel of BC (box counting) was used after setting "Num G" to 4 in the grid design settings. Under graphical options, check the metrics box to analyze the cell's convex hull and bounding circle. After the settings are complete, click on SC (scan). The $D_B$ of microglia can be viewed as the average rate of change in detail with change in resolution sampled from the image.

**Microglia volume and size measurements**

IMARIS measured microglia's surface area and volume after rendering a SIM image of the microglia via the surface panel.


μm

### Extended Figure Legends

#### Fig. S1. Microglia status in the fetal brains with different gw

**A.** The whole brain stitching images of CD34 and IBA-1 antibodies immunostaining in the fetal brain with 7.5, 9, and 15 gw. PAX6 and OTX2 antibodies immunostaining for outlining the forebrain and midbrain (the left in the lower panel). Single-channel images of IBA-1 immunostaining are shown as grey. The right panel is the magnified image of the boxed region. IBA-1$^{+}$ cells are predominantly localized in the intermediate zone (IZ.) and subplate (SP), with a limited presence in the subventricular zone (SVZ) and a few in the mantel zone/cortical plate (MZ/CP) region.

**B.** Microglia density in the fetal brains with 7.5, 9, and 15 gw. Fiji ImageJ did the counting.

**C.** The 3D ultrahigh-resolution images of SIM microscopy revealed microglia's morphological shifts during increasing gw.

**D.** The structural analysis of microglia in different gw shows the increase of microglia's complexity following gw.

#### Fig. S2. The whole brain immunostaining image with IBA-1 and Ki67 antibodies in the mouse fetal brain at 12.5 days by the CUBIC method

**A.** A large proliferative microglia aggregate stained by IBA-1 and Ki67 antibodies in the fetal brain with 13 gw (The dashed line, the margin of caudate).

**B.** 3D whole view of mouse whole head immunostained with IBA-1 and Ki67 antibodies. The staining was performed using the CUBIC method, and Nikon A1 was scanned with a 10X objective.

**C.** Three representative sectional views of the whole brain scanning images revealed that large microglia aggregates with high proliferative potential did not exist in the mouse fetal brain at 12.5 days (right corner, the navigation).

**Fig. S3. The cross-talking pattern of SV40 microglia with the induced striatal neurons in 2D coculture**

**A**. Schematic illustration of getting induced-striatal neurons from ES or hiPSC.

**B.** Ki67, Sox2 and Tuj-1 or DARPP32 and CTIP2 or GABA and MEIS1 antibodies immunostaining of 2D neurons revealed that a significant amount of DARPP32$^+$ and GABA$^+$ neurons exist in the induced system.

**C.** The representative images of induced neurons and SV40 microglia in coculture revealed that SV40 microglia conjugated with or penetrated the neuronal fibers at day 41 and day 42 (yellow arrows, penetrated microglia); The right panels are a larger view that reveals the full contact of microglia with induced neurons. The upper panel is a phase contract microscopy image, and the lower panel shows the status of microglia.

**D.** The morphology of SV40 microglia that cocultured within 2D neurons for 24h revealed a bipolar morphology of microglia with longer processes, and the neurons without neurons mostly exhibited a plated morphology. The bar plots of the length of projections and endpoints on microglia revealed that the complexity of SV40 microglia increased in coculture (Endpoints represent all protrusions on the microglia cytosol, with no distinction between primary or secondary protrusions). Data, mean ± S.E.M; *One-way ANOVA*; *, $p < 0.05$, **, $p < 0.01$; ns, no significant difference.

**E, F.** Ki67, BDNF, and IBA-1 immunostaining revealed that SV40 microglia nested in neuronal fibers and expressed BDNF and Ki67.

**G.** SIM images of penetrated SV40 microglia in the deep region of hCO stained by GFP, PSD95, and MAP2 antibodies revealed PSD95$^+$ and MAP2$^+$ puncta in SV40 microglia. IMARIS does the rendering.

**Fig. S4. Introducing SV40 microglia into Day 60 hCO**

**A.** The morphology of SV40 microglia recruited to the surface of Day 60 hCO.

**B-D.** The immunostaining images of PAX6 and GFP antibodies (B) or GFP, Tuj-1, and CTIP2 (C) or Ki67, pH3, and GFP antibodies (D) in hCO with SV40 microglia that cocultured for 7 days. The bar plots are the count of PAX6$^+$ and CTIP2$^+$ cells in hCO with or without SV40 microglia (yellow arrows, penetrated microglia; the boxed region, the magnified region in the right panel). Data, mean± SD; t-test. Repeat: n = 4. Organoids in each group: n = 12

**E.** Comparing the morphology of penetrated SV40 microglia in hCO with human microglia in the SVZ region of 9 gw fetal brain. The penetrated SV40 microglia in SVZ displayed a bipolar morphology that is identical to the microglia in SVZ of the fetal brain with 9.5 gw (yellow arrows, penetrated microglia).

**Fig. S5. The characteristics of microglia in DS brain regions**

**A**. A proliferative large microglia aggregate in DS brain immunostained with Ki67 and IBA-1 antibodies (the boxed region in the left panel, the magnified region in the right panel).

**B.** The images of RG scaffolding microglia in the DS brain revealed that RG scaffolding microglia harbor multiple complicated processes (the boxed region, the magnified image in the right panel).

**C.** Comparing count of the processes in RG scaffolding microglia in DS brain with a healthy fetal brain. Data, mean ± s.e.m.

**D.** SIM images revealed the characteristics of processes and bulbs of microglia in the CP of the DS brain. A microglia with observable thick processes (i); a microglia with multiple bulbs on the stem of processes in the CP region (ii); a Ki67$^+$ microglia with enormous processes and blubs in CP region (iii) (white arrows, enlarged stem; yellow arrows. bulbous ending; pink arrows, stem bulbs).

**Fig. S6. The apoptotic and immunological characteristics of DS brain**

**A.** SIM images of DS and healthy fetal brain immunostained by caspase-3 and IBA-1 antibodies revealed that DS brain contains caspase 3$^+$ large projection neurons with fragmented nucleus (i) or caspase 3$^+$ multiple torturous processes (ii) (white arrows, caspase 3$^+$ cells or fragments). Caspase 3$^+$ cells or processes are rare in the microglia aggregate in healthy and DS brains (lower panels).

**B**. Comparing the area fraction of caspase 3$^+$ region in DS fetal brain with healthy fetal brain.

**C.** The stitched 3D images of IBA-1, CD8, and CD177 antibodies immunostaining revealed that CD8$^+$ T-cells and CD177$^+$ neutrophils are absent in the region of microglia aggregates in both healthy and DS fetal brains (the inner insert in LGE panels, the typical CD8$^+$ T cell in microvessels).

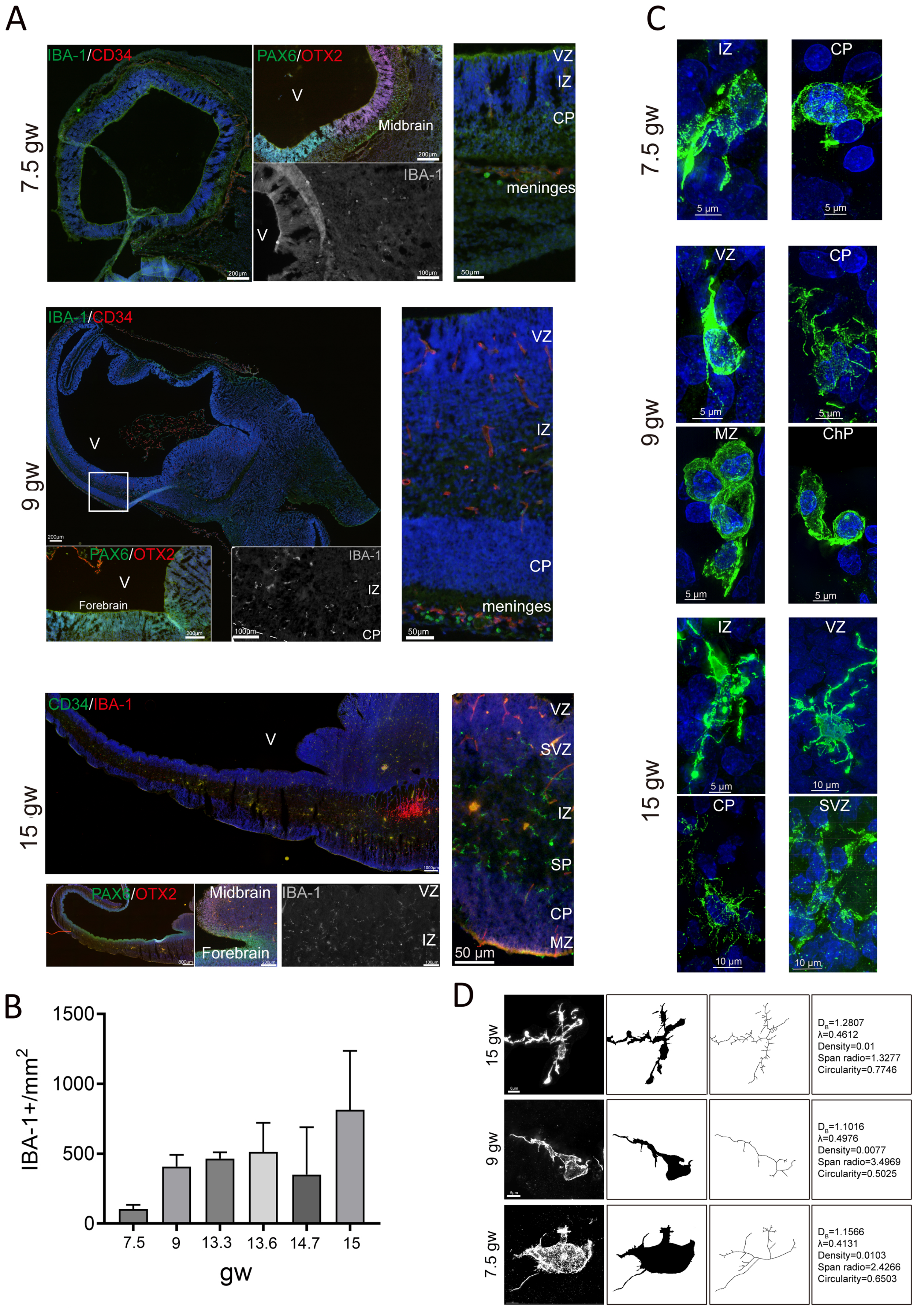


Fig. S1 Song et al.

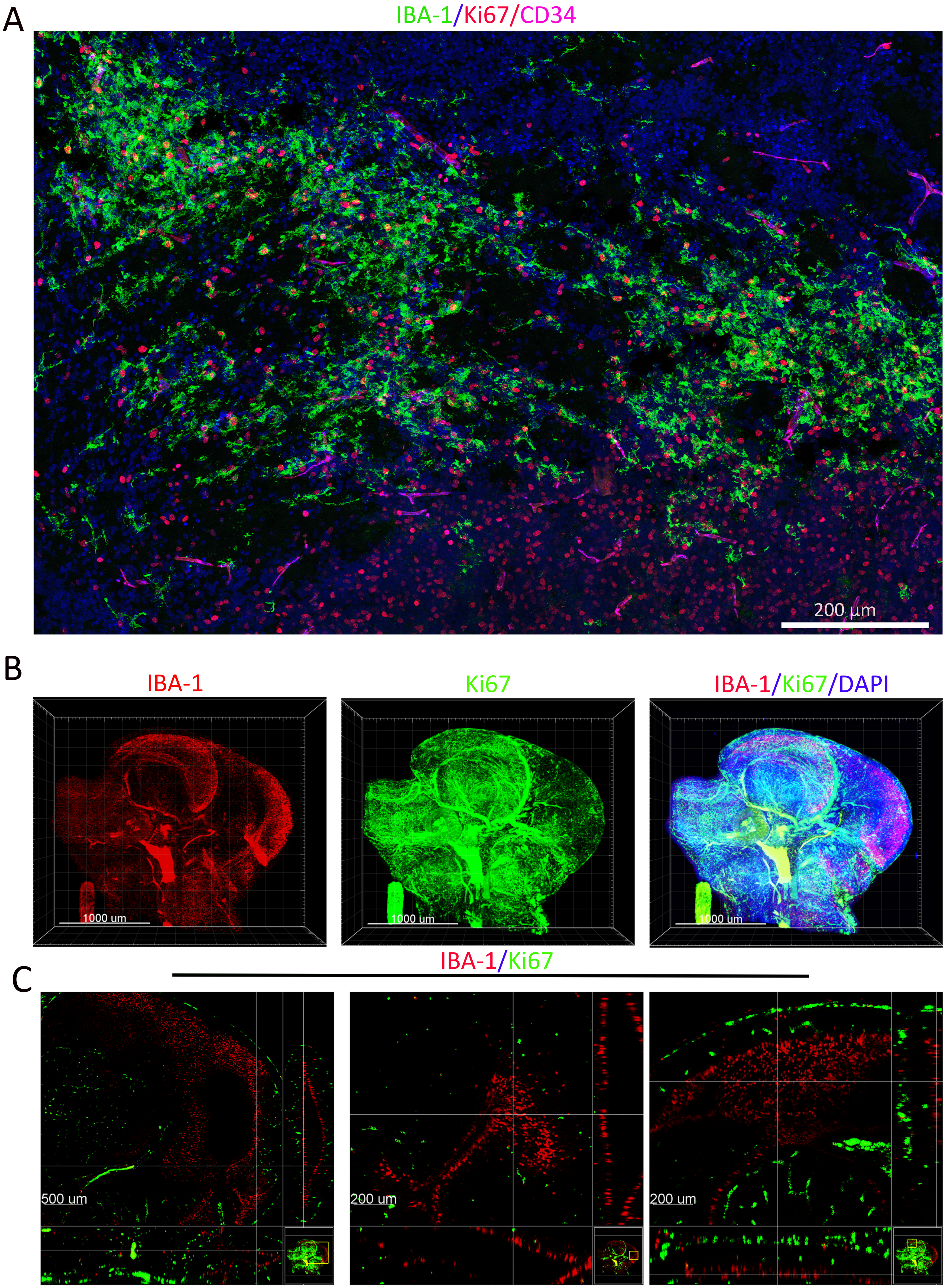


Fig. S2 Song et al.

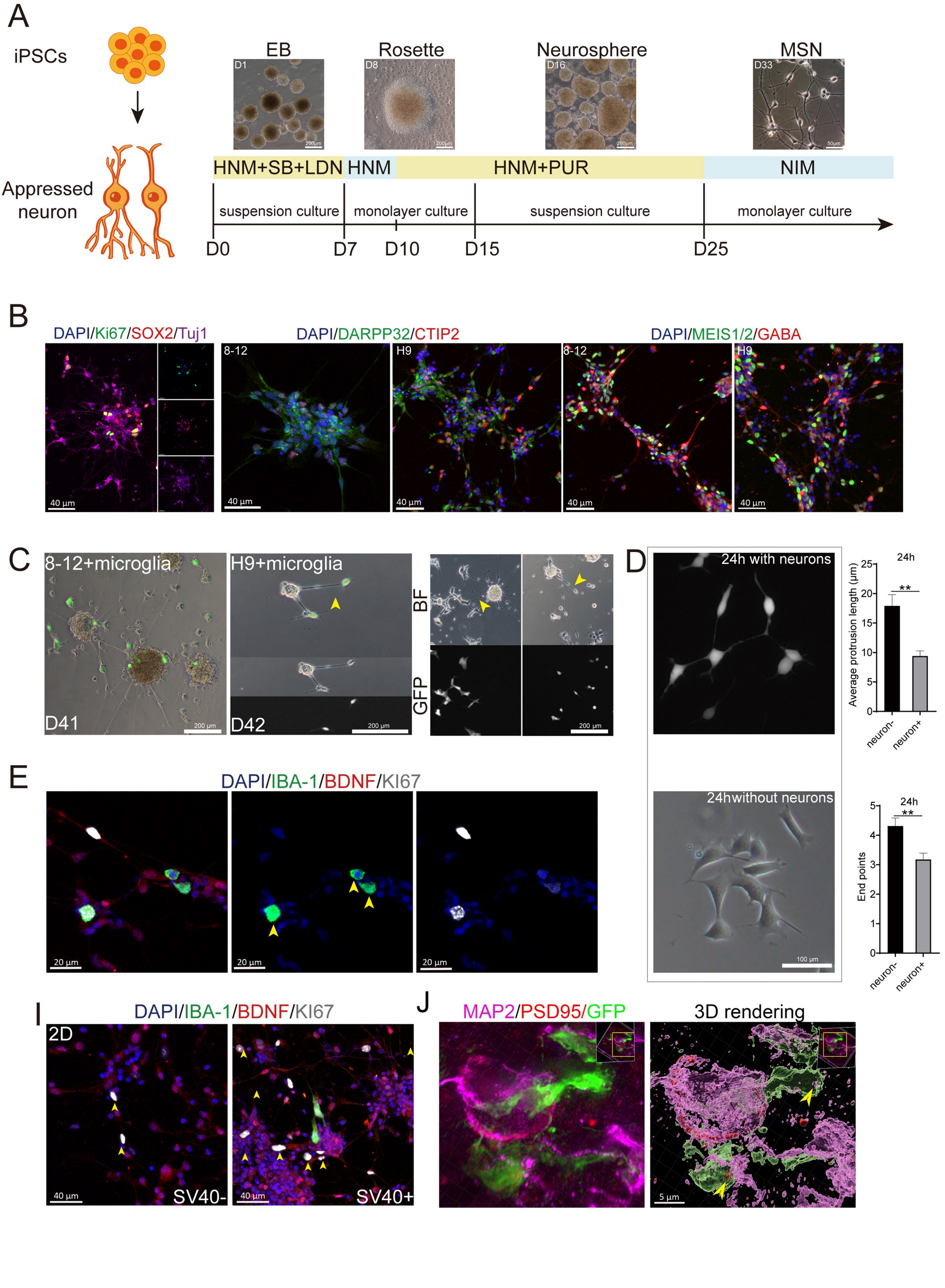


Fig. S3 Song et al.

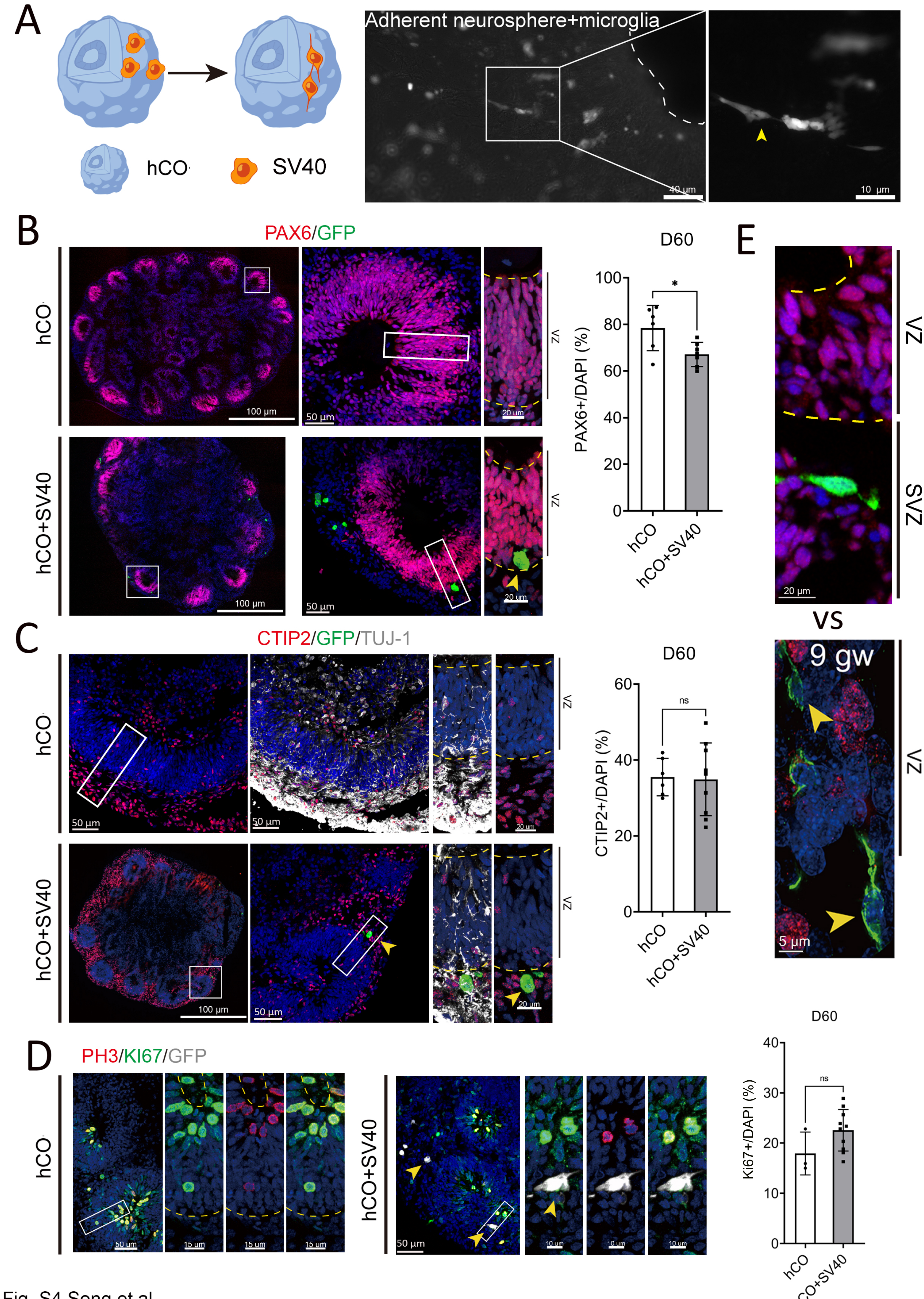


Fig. S4 Song et al.

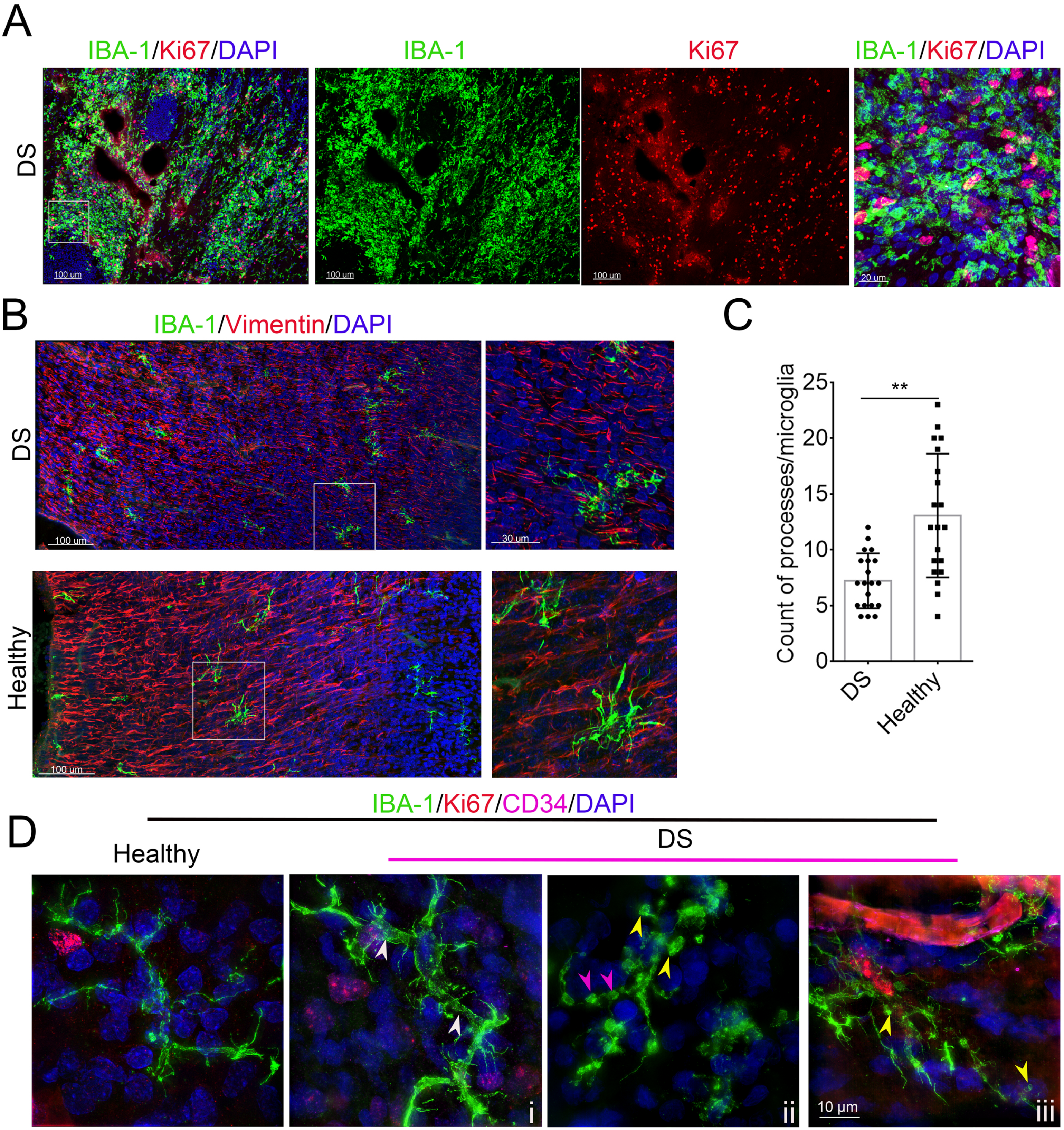


Fig. S5 Song et al.

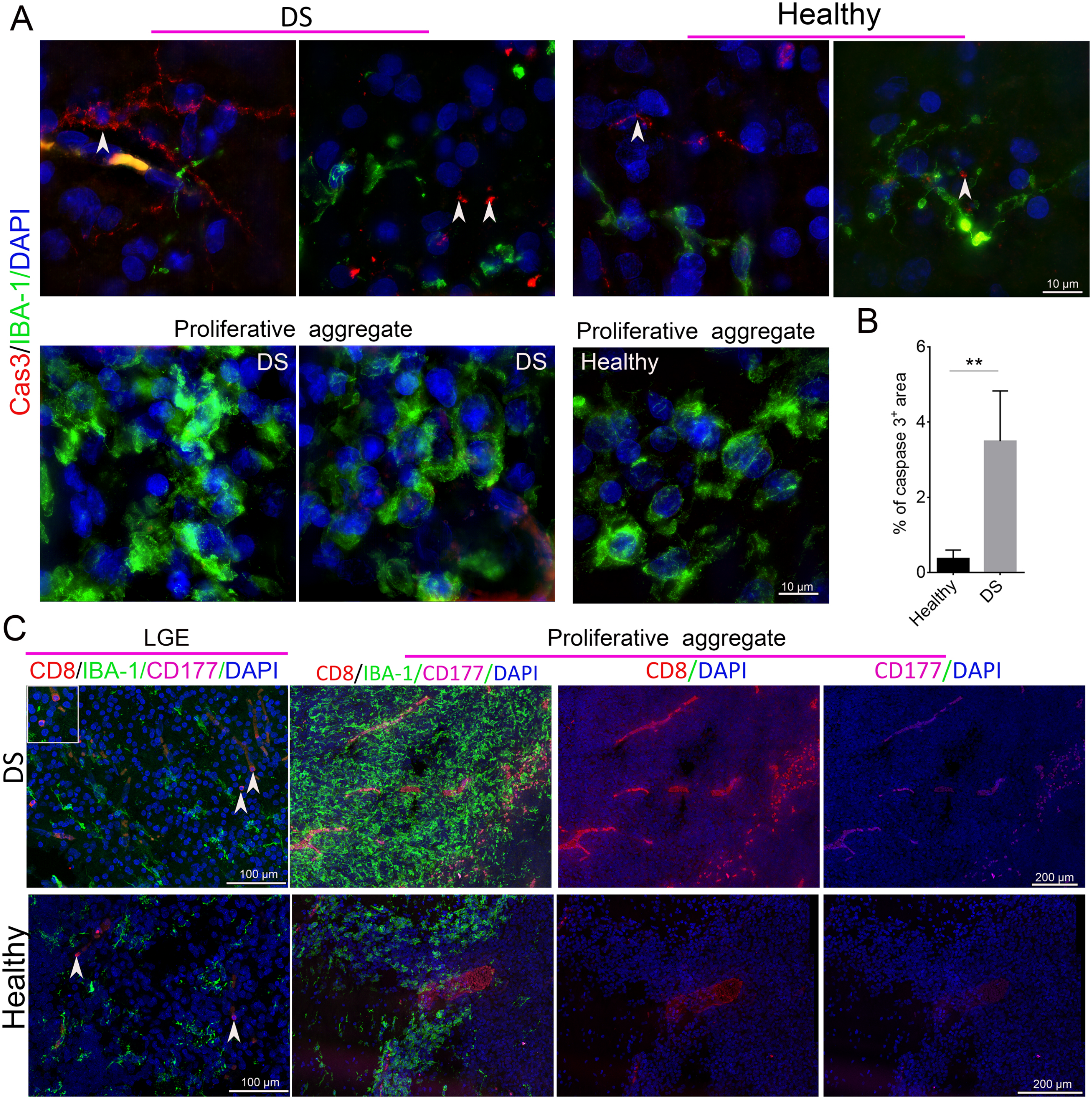


Fig. S6 Song et al.

Table S1. Summary of the proliferative microglia aggregates in fetal brains

| NO. | Gestation week | Proliferate center ($mm^2$) | Total IBA-$1^+$ cell counted | Total Ki67$^+$ cell counted | Ki67$^+$/IBA-1 Average (%) | Cerebral hemisphere length (cm) |
|---|---|---|---|---|---|---|
| FBr-1 | 13 gw+4 days | 0.108 | 3,727 | 535 | 14.355 | 2.532 |
| FBr-2 | 13 gw+2 days | 1.667 | 1725 | 430 | 24.928 | 2.876 |
| FBr-3 | 15 gw | 1.511 | 2,904 | 853 | 29.37 | 2.393 |
| FBr-4 | 14 gw+5 days | 0.962 | 115 | 13 | 11.304 | 3.514 |
| FBr-5 | 16 gw | 2.129 | 947 | 191 | 20.168 | N.A |
| FBr-6 | 15 gw | 0.577 | 474 | 177 | 37.341 | 3.594 |
| FBr-7 | 13 gw+4 days | 0.795 | 1142 | 393 | 34.441 | 3.639 |
| FBr-8 (DS) | 16 gw+3 days | 4.168 | 4016 | 1141 | 28.411 | 3.860 |

*FBr, Fetal Brain; DS, Down's syndrome. Each FBr section ≥ 3.
